# Label-free Assessment of Complement-Dependent Cytotoxicity of Therapeutic Antibodies via a Whole-Cell MALDI Mass Spectrometry Bioassay

**DOI:** 10.1101/2024.05.03.592336

**Authors:** Stefan Schmidt, Alexander Geisel, Thomas Enzlein, Björn C. Fröhlich, Louise Pritchett, Melanie Verneret, Christian Graf, Carsten Hopf

## Abstract

Potency assessment of monoclonal antibodies or corresponding biosimilars in cell-based assays is an essential prerequisite in biopharmaceutical research and development. However, cellular bioassays are still subject to limitations in sample throughput, speed, and often need costly reagents or labels as they are based on an indirect readout by luminescence or fluorescence. In contrast, whole-cell Matrix-Assisted Laser Desorption/Ionization Time-of-Flight (MALDI-TOF) Mass Spectrometry (MS) has emerged as a direct, fast and label-free technology for functional drug screening being able to unravel the molecular complexity of cellular response to pharmaceutical reagents. However, this approach has not yet been used for cellular testing of biologicals. In this study, we have conceived, developed and bench-marked a label-free MALDI-MS based cell bioassay workflow for the functional assessment of complement-dependent cytotoxicity (CDC) of Rituximab antibody. By computational evaluation of response profiles followed by subsequent *m/z* feature annotation via fragmentation analysis and trapped ion mobility MS, we identified adenosine triphosphate and glutathione as readily MS-assessable metabolite markers for CDC and demonstrate that robust concentration-response characteristics can be obtained by MALDI-TOF MS. Statistical assay performance indicators suggest that whole-cell MALDI-TOF MS could complement the toolbox for functional cellular testing of biopharmaceuticals.

## Introduction

Discovery and development of biopharmaceuticals such as therapeutic antibodies requires extensive developability assessment in suitable bioassays that address safety (e.g. immunogenicity), physicochemical properties (e.g. aggregation), formulability and biological activity^1-4^. The comparison of potencies of different technical or production antibody batches or of biosimilars to their antibody reference products requires bioassays that address a wide range of mechanisms of action (MoA) such as antibody-dependent cell-meditated cytotoxicity (ADCC), antibody-dependent cellular phagocytosis (ADCP) or complement-dependent cytotoxicity (CDC), and that are robust and amenable to thorough statistical analysis^5-7^. In CDC bioassays, antibodies like the anti-CD20 therapeutic Rituximab bind to their antigens on target cells like human B cells or B lymphoblastoid Raji cells derived from a patient with Burkitt lymphoma. Via their Fc regions, they recruit the complement system, which in turn opsonizes and lyses the target cells via the formation of a membrane attack complex^8^.

In recent years, radioactive techniques for CDC measurement such as the ^51^Cr release method have been replaced by non-radioactive approaches like flow cytometry, fluorescence or luminescence measurements^9-11^. Nevertheless, these techniques still require labels and/or often costly reagents. In contrast, mass spectrometry (MS)-based assay technologies based on Matrix-Assisted Laser Desorption/Ionization (MALDI) or on acoustic mist ionization/acoustic ejection are label-free, and offer nowadays the necessary speed and analytical depth for information-rich, high-throughput compound profiling and screening^12-16^. Since the inception of MALDI MS-based cellular potency bioassays by Munteanu et al.^17^ several applications relating to mechanistic effects on cellular targets, phenotypic assessment of MoA, compound uptake and its inhibition, stem cell differentiation, glucose-stimulated insulin secretion and more have been described^15,16,18-22^. However, no implementation of MS-based cell assays in the field of biologics research and development has been reported so far.

In this study, we have conceived, designed and developed an initially untargeted MALDI-TOF MS-based CDC bioassay for Raji cells and benchmark it against a common luminescence assay. We employ tandem-MS fragmentation and ion mobility analysis on a timsTOF mass spectrometer to identify adenosine triphosphate (ATP) and glutathione (GSH) as suitable response markers employed in a targeted MALDI-TOF MS CDC bioassay. Finally, we introduce new statistical measures for quality assurance in MALDI MS-based assays.

## Results and Discussion

### Experimental workflow of the whole-cell MALDI-TOF MS CDC bioassay

Leveraging recent strategies for automated MALDI-TOF MS-based cell assays for small-molecule drug-like compounds^14,15^, we aimed to develop a MALDI-MS cell assay for the label-free potency assessment of therapeutic antibodies. As a proof-of-concept, we chose a standard biopharmaceutical potency bioassay that measures complement-dependent cytotoxicity (CDC) of the monoclonal antibody Rituximab (anti-CD20)^23^ The classical implementation of the CDC bioassay involves incubation of Rituximab samples in different concentrations with rabbit complement and target cells (Raji cells) in microtiter plates, in order to induce complement activation, formation of a membrane-attack complex and finally cell lysis. Target cell killing is then measured by monitoring cell viability utilizing a luminescence-based reagent kit and a microplate reader, followed by statistical data analysis to obtain concentration response characteristics like the logarithmic value of the half maximal effective concentration pEC_50_.

For the implementation of the MALDI-MS assay, it was necessary to optimize the CDC reaction conditions, e.g. number of cells, to ensure information-rich and robust MS readout. To this end, we conceived and developed a (semi-)automated MALDI-MS assay protocol based on published procedures^15,19^ consisting of two main parts: i) complement activation and sample plate preparation using a liquid handling robot and ii) metabolic, whole-cell MALDI-TOF-MS readout and subsequent multivariate statistical data analysis. Raji cells were treated in 96-well plates with complement at a 1:2 dilution and Rituximab in a concentration range of 7.3 ng/mL to 0.12 mg/mL for 120 min at 37 °C to induce CDC and cell lysis. The luminescent CDC assay was performed in parallel to act as a comparator (**Fig. 1**). To account for possible MALDI matrix suppression effects, a suitable cell number is essential^15,19^. Raji cells were harvested for 24h to 48h (**Fig. S1a**) and suspended at 5.0·10^6^ cells/mL for an optimal number of 5000 cells per MALDI target spot (**Fig. S1b**). Complement and antibody concentrations were both optimized to yield a fold-change in cell viability of about 5-fold or more (**Fig. S1c**) between cells treated with complement only (negative control), and cells treated with both complement and mAb. Viability of complement/mAb-treated cells was determined relative to non-treated cells. In addition, cells treated with Triton X-100 to cause cell death and lysis served as positive controls.

**Figure 1.**
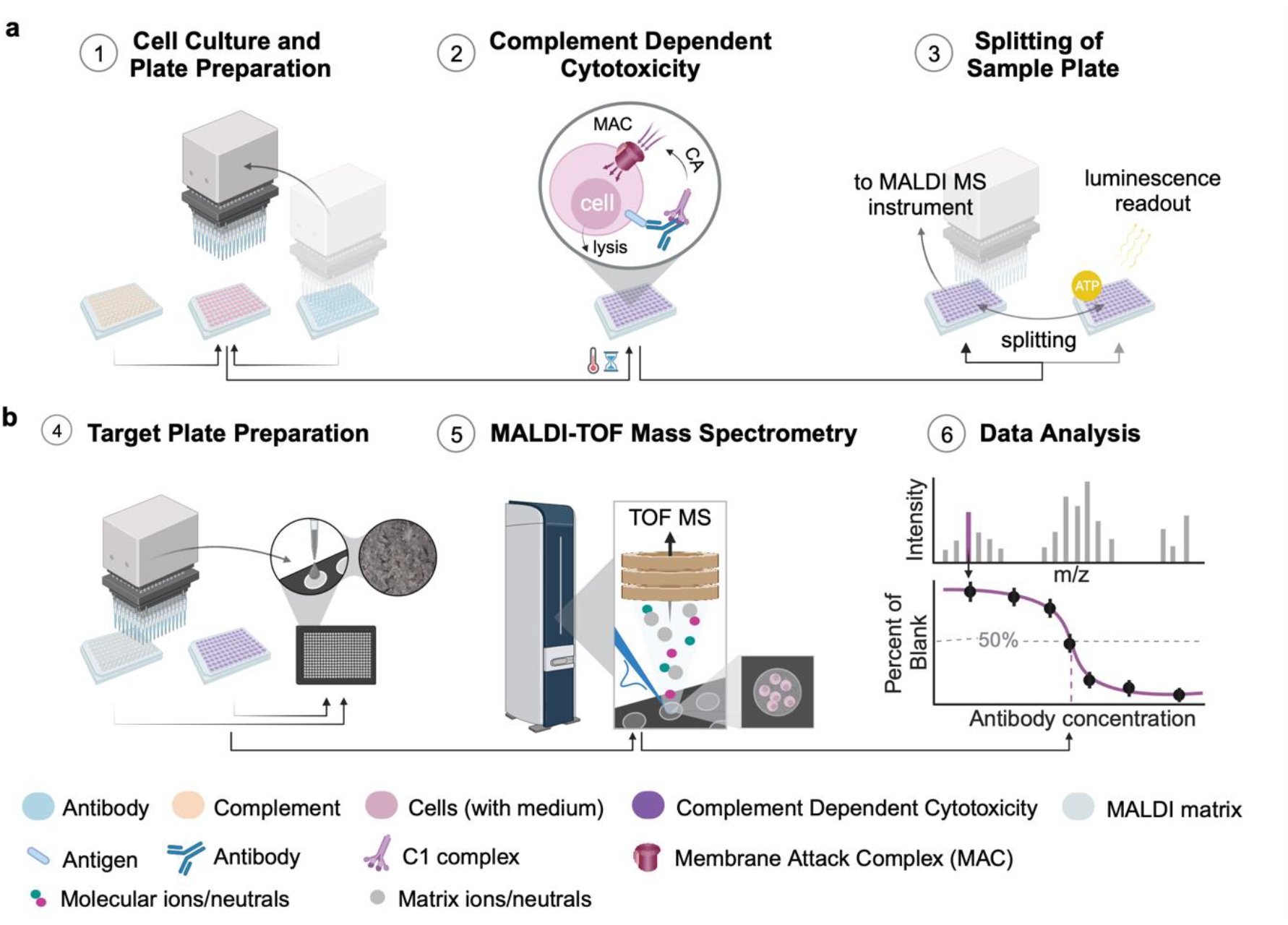
Schematic overview of the MALDI MS-based complement-dependent cytotoxicity (CDC) cell assay. **a**, Complement activation (CA) and sample plate preparation: (1) Raji cells are seeded in 96-well assay plates. (2) Rabbit complement and Rituximab dilutions are added by an automated liquid handling system to ensure high reproducibility. During subsequent incubation of the resulting assay plate at 37 °C, 5% CO_2_ for 2h, complement is activated by cell-bound monoclonal antibodies (mAbs) and forms the membrane attack complex that triggers cell lysis. (3) Thereafter, a small sample is analyzed in a luminescence viability assay as reference readout. **b**, Molecular readout by rapid MALDI-MS. (4) The remaining sample is processed for MALDI-MS readout. This includes centrifugation-, washing- and freezing steps as well as resuspension of cells in solvent. Subsequently, the sample is automatically spotted onto a MALDI target plate (*n*=4 measurement replicates per well). MALDI matrix (with spiked-in stable isotope labeled internal standard) is deposited by either spotting or spraying. (5) Mass spectra are acquired using rapifleX or timsTOFflex mass spectrometers. (6) Multivariate statistical analysis reveals response markers, which are further used to determine EC_50_ values for the tested antibodies. Created with BioRender.com.

High repeatability and careful sample handling were ensured by an automated liquid handling system. The number of cells left after resuspension and washing was consistently around 55% (**Fig. S1d**) for all antibody concentrations. Four wells of cell were prepared per treatment condition to assess inter-well variability. Cells from each well were split onto four target spots on 384-well MALDI target using laboratory automation to assess spot-to-spot variability. Transferred cells were embedded in MALDI matrix that was spiked with an internal standard for normalization in positive ion mode. Measurements were repeated in triplicates on different days.

### Untargeted identification of metabolic CDC markers by MALDI MS

MALDI-MS spectra of complement/Rituximab-treated Raji cells were information-rich in both positive and negative ion mode, typically featuring 400 to 500 *m/z* values that were peak-picked (signal-to-noise (S/N) > 3) (**Fig. 2a, i**). We noted striking differences in the m/z fingerprints between treated and untreated cells indicating putative concentration-response and, hence, potential CDC markers (**Fig. 2a, ii**). For automatic, untargeted identification of metabolic response markers^14^, spectral data in positive and negative ion mode were analyzed independently by their concentration-response characteristics.

**Figure 2.**
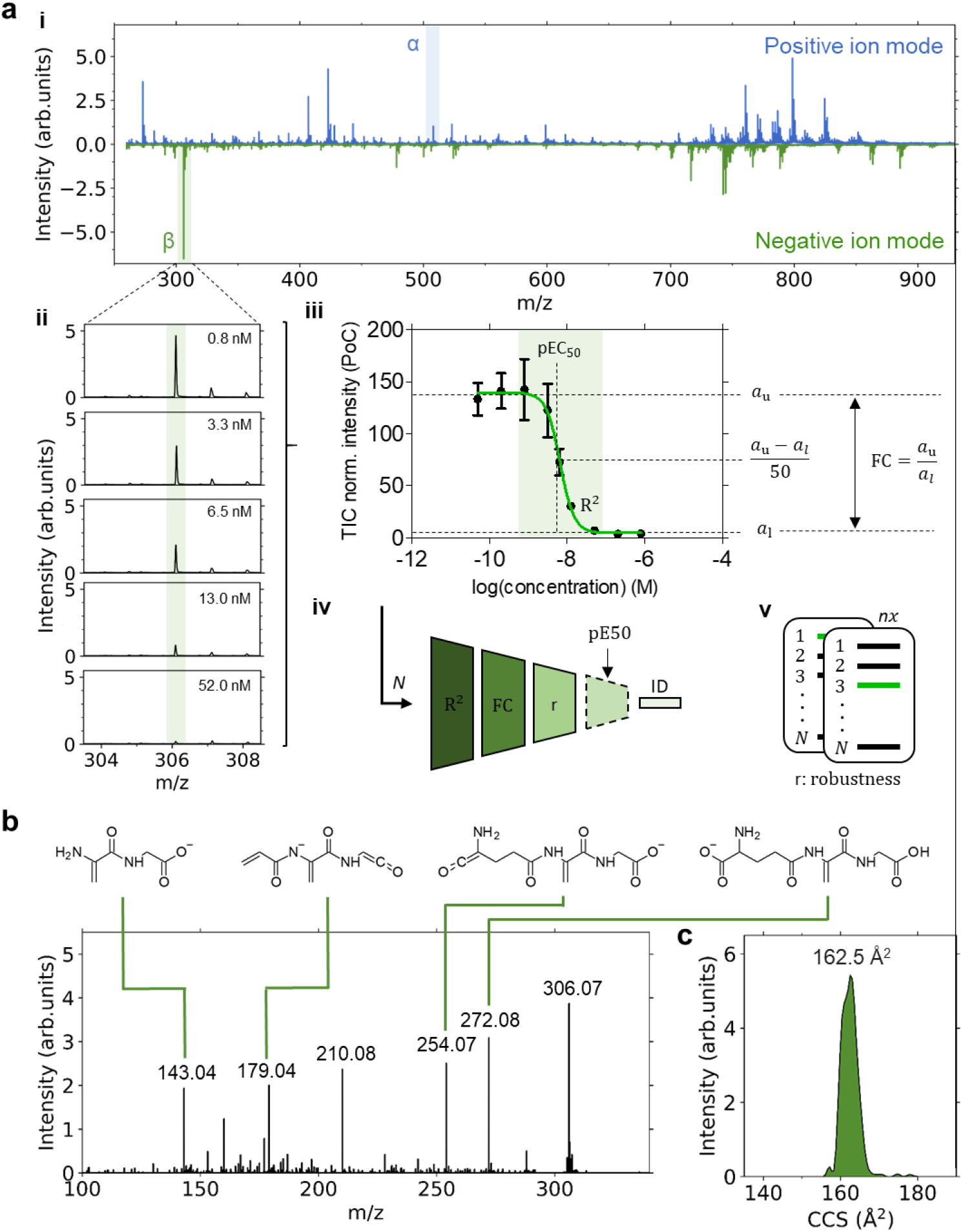
Untargeted identification and validation of metabolic CDC response markers by MALDI TOF MS. **a**, Mass spectra for positive and negative ion mode (i) are independently automatically screened for potent marker candidates that exhibit a concentration-dependent response (ii and iii) as exemplarily shown for the molecule at *m/z* 306.08 (β). After an initial peak-picking step, for each *m/z* feature, a concentration-response curve is fitted to the data. Candidates are judged and filtered (iv) based on their respective value for the coefficient of determination (R_2_), the fold-change (FC) defined by the ratio of the upper and lower asymptotes, *a*_u_ and *a*_l_, their robustness, i.e. quality of the data for serval replicates *n* (v), and optionally by their pEC_50_ value in comparison to a known reference value. Finally, the most promising marker(s) is (are) annotated by its (their) unique fragment pattern (**b**) and/or their *m/z* and CCS value (**c**) as demonstrated for glutathione (theoretical *m/z* 306.0765 for [M-H]^-^; NEDC matrix). MALDI-MS/MS spectra of *m/z* 306.08 display unique fragments of glutathione as indicated by corresponding fragment structures. Measurements were performed on the timsTOF flex mass spectrometer with a collisional energy of 18 eV. **c**, Ion mobilogram of a “blank spectrum”, i.e. non-treated cells, for experimental deduction of collisional cross section (CCS) of 163 ± 2 Å^2^ for glutathione.

Data fitting using a 4-parameter logistic regression model (**Fig. 2a, iii**) enabled potential *m/z* candidates to be filtered and ranked according to their coefficient of determination (R^2^), their fold-change (FC), i.e. ratio between the values for upper and lower asymptotes *a*_u_ and *a*_l_ of the fit (**Fig. 2a, iv**), and their robustness/repeatability. Optionally, the respective pEC_50_ values may be compared against a known reference value. Here, we used results from the reference luminescence assay as a benchmark. Based on this feature selection process, the most prominent and reliable *m/z* candidate for each polarity, *m/z* 306.08 and *m/z* 508.00, was selected and identified by its MS/MS spectrum and ion mobility as glutathione and adenosine triphosphate, respectively (**Fig. 2b and S3a**). Both molecules are present at millimolar concentrations in cells and are among the most abundant intracellular metabolites^24-26^.ATP release is measured in the CDC ATP-based cell viability luciferase reference assay.

To investigate the presence of unrelated compounds or molecular entities, e.g. MALDI matrix clusters interfering with the identified markers that are within the mass resolution of the MS instrument, ion mobility spectrometry was performed. Our experimentally deduced collisional cross section (CCS) values of 198 ± 2 Å^2^ (ATP), and GSH, 163 ± 2 Å^2^ (GSH) are comparable with CCS values predicted by machine learning approaches^27,28^ and that are listed in the Human Metabolome Database (HMDB; http://www.hmdb.ca) [196.8 Å^2^ for ATP (HMDB0000538) and 161.9 Å^2^ for GSH (HMDB0000125)(**Fig. 2c and S3b**). The presence of interfering additional *m/z* features is therefore unlikely. Both markers displayed clear concentration-responses and were readily discernible when analyzed by faster MALDI-TOF MS (**Fig. 3**). In case of ATP, an internal standard (^13^C_10_, ^15^N_5_) was used for normalization, whereas for GSH we relied on normalization on the total ion current (TIC).

**Figure 3.**
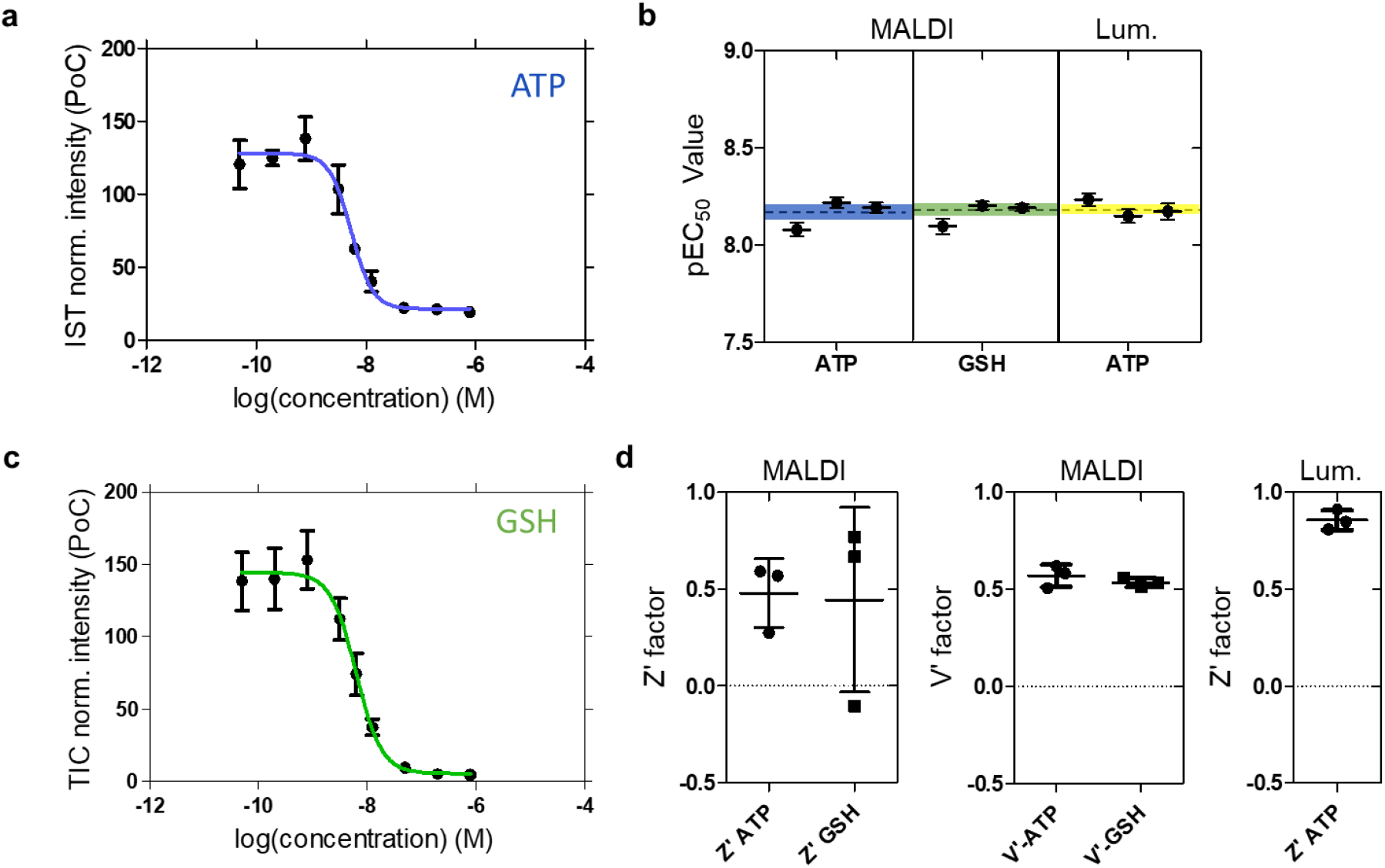
Characteristics of the MALDI MS-based CDC cell assay for reference Rituximab sample #ref. **a**, Concentration-response curve for ATP ([M+H]^+^, *m/z* 508.02). Signal intensities were normalized to the internal standard (IST; as percent of control (PoC). **b**, pEC_50_ values for three biological replicates. The mean and standard deviation are indicated by the dashed line and color-filled areas, respectively. pEC_50_ values were 8.2 (N=4), 8.2 (N=2) and 8.2 (N=2) for the MALDI MS assay using ATP and GSH as well as for the luminescence (lum.) ATP reference assay, respectively. **c**, Concentration-response curve for glutathione ([M-H]^-^, *m/z* 306.08) after TIC normalization. In **a** and **c**, each data point is defined by the mean and standard deviation of three biological replicates. **d**, Assay quality measures, values for the *Z*‘and *V*’ factors, for the individual assay conditions. Mean and standard deviation across three biological replicates are indicated.

### Method capability and assay performance

In order to assess the MALDI MS method capability to determine the relative potency, i.e. pEC_50_ values, of different Rituximab batches, we performed a comparability analysis between different Rituximab drug product batches with #ref as reference (*n*=3; **Tab. S1**). The potency of each antibody was determined by fitting a 4-parameter logistic regression model to the metabolic response data to obtain pEC_50_ values (**Fig. 3a to 3c**). To simultaneously evaluate results for all antibodies and both metabolic readouts (ATP and GSH), in total 24 response curves (*n*=3, 2(+2) mAbs (and reference), 2 marker molecules) were measured (**Fig. S3-S4**). Then, pEC_50_ values were benchmarked against the corresponding luminescence reference assays (**Tab. S2**; **Fig. 3b** for reference antibody #ref as example), yielding mean pEC_50_ values of 8.16 ± 0.04 (ATP), 8.18 ± 0.02 (GSH) and 8.19 ± 0.02 (luminescence assay). For all antibodies batches measured, the mean pEC_50_ values from metabolic ATP and GSH MALDI-TOF MS readouts were highly consistent with the luminescence results (**Tab. 1 and S1)**. Evaluation of the calculated relative pEC_50_ values of the three different commercial Rituximab batches **(Tab. 2)** showed that the MALDI MS-based readout provides values with acceptable accuracy and variability, which compares well with the results from the luminescence method.

**Table 1.** pEC_50_ values for CDC activity of different Rituximab (Mabthera**®**) batches (#ref as reference) and readouts. ^1^Mean pEC_50_ value and their uncertainty for three biological replicates (see Table S2 for pEC_50_ values of each technical replicate). ^2^Mean pEC_50_ values of all mean values for a given antibody.

| Type of readout | Measure | #ref | #1 | #ref | #2 |
| --- | --- | --- | --- | --- | --- |
| MALDI ATP | pEC<sub>50</sub><sup>1</sup> | $8.18 \pm 0.06$ | $8.14 \pm 0.03$ | $8.16 \pm 0.04$ | $8.09 \pm 0.03$ |
| MALDI GSH | pEC<sub>50</sub><sup>1</sup> | $8.19 \pm 0.01$ | $8.20 \pm 0.02$ | $8.18 \pm 0.02$ | $8.18 \pm 0.01$ |
| Luminescence | pEC<sub>50</sub><sup>1</sup> | $8.18 \pm 0.04$ | $8.21 \pm 0.05$ | $8.20 \pm 0.03$ | $8.18 \pm 0.03$ |
| | pEC<sub>50</sub><sup>2</sup> | $8.19 \pm 0.01$ | $8.18 \pm 0.02$ | $8.19 \pm 0.01$ | $8.17 \pm 0.01$ |

**Table 2.** Relative pEC_50_ values for CDC activity of two Rituximab (Mabthera**®**) batches (#1 and #2) in comparison to the reference batch (#ref). ^1^Results obtained from the luminescence readout performed in parallel to the MALDI readout.

| Type of readout | Measure | #ref | #1 | #2 |
| --- | --- | --- | --- | --- |
| MALDI ATP | Relative pEC<sub>50</sub><br>(%) | 100 | $110 \pm 17$ | $117 \pm 13$ |
| MALDI GSH | Relative pEC<sub>50</sub><br>(%) | 100 | $98 \pm 5$ | $100 \pm 5$ |
| Luminescence<sup>1</sup> | Relative pEC<sub>50</sub><br>(%) | 100 | $93 \pm 14$ | $105 \pm 10$ |

To further evaluate MALDI-TOF MS assay quality, the two independent measures Z’- and V’-factors^29,30^ for assay robustness were determined for the reference antibody #ref (**Fig. 3d)**. Except for one replicate with high variability, Z’-values were between 0.5 and 0.6 for the ATP readout and between 0.6 and 0.8 for the GSH readout. Z’ values for the luminescence assay were above 0.8. The fact that all V’ values are found to be within 0.5 and 0.65 for both metabolic readouts indicated that even the off-replicate with high variance provided acceptable determination of potency (**Fig. 3b)**.

In general, the sources of variation within data in MALDI MS-based bioassays are typically unknown and could originate from either sample preparation steps (**Fig. 1a**) like cell washing/ resuspension or from the MALDI-TOF MS readout itself (**Fig. 1b**), including target plate preparation steps. To statistically assess the main source of variation, we investigated the inter-well (four cell culture wells per plate) and spot-to-spot (four MALDI target spots per cell culture well) variability of the bioassay. To this end, we introduced a statistical measure 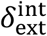 (**Fig. 4a**) with the following meaning: for 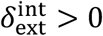 the spot-to-spot variability is larger than the inter-well variability. Accordingly, for 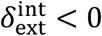 it is smaller than the inter-well variability. First, we determined the mean and its uncertainty (**Fig. 4b**) for each of the four spot-to-spot measurement replicates per well and individual treatment condition (i-v in **Fig. 4b**) to further deduce the internal *δ*_int_ and external *δ*_ext_ errors for each treatment condition from which 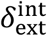 is calculated. Internal and external errors are commonly used as estimates for the uncertainty within the calculation of the mean to test the statistically consistency within data^31^.

**Figure 4.**
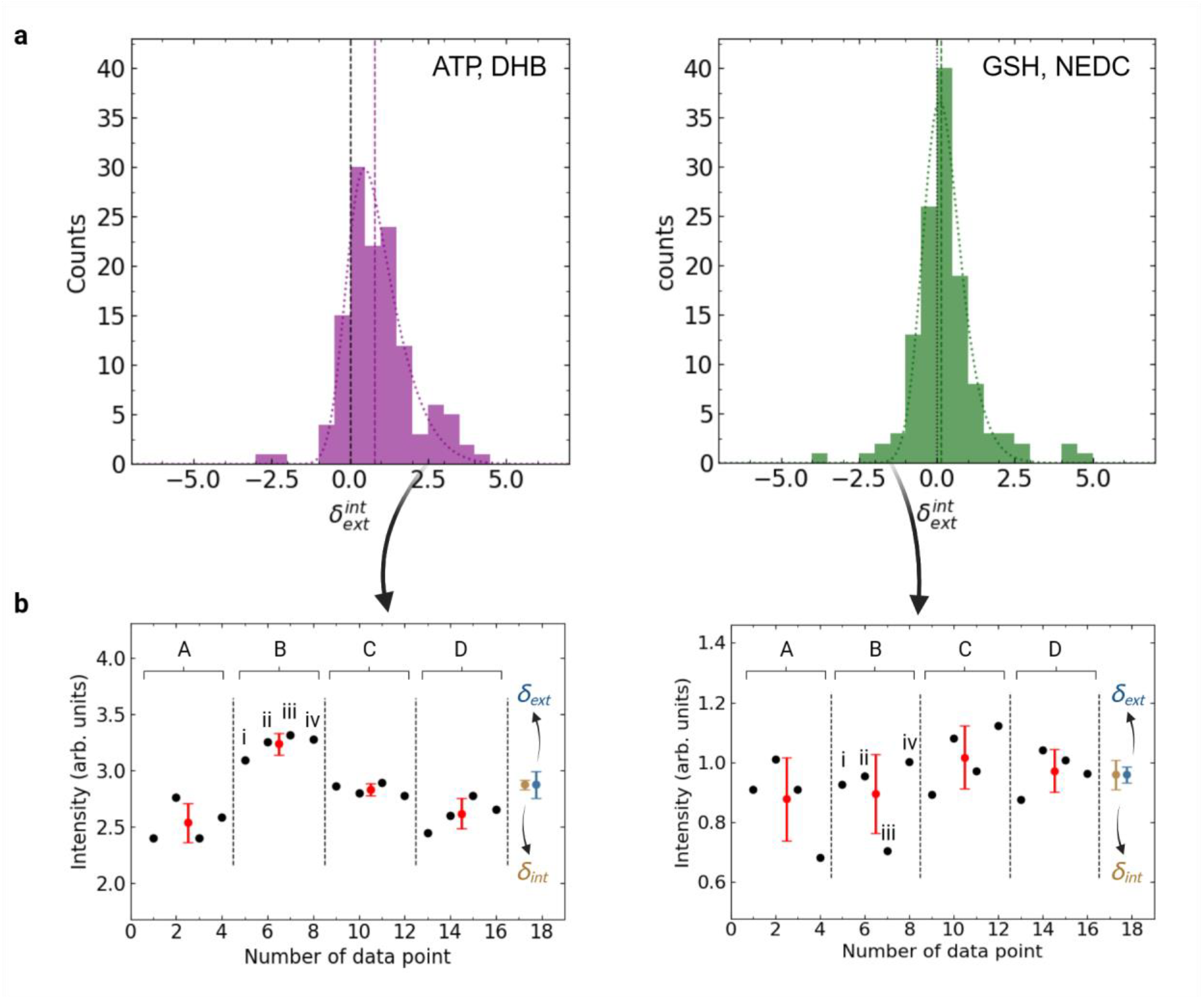
Comparison of inter-well and spot-to-spot variability. **a**, Histogram of modified ratio of the external and internal error 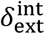 (see method section) for two different assay conditions, MALDI MS ATP readout using DHB matrix and MALDI MS GSH readout using NEDC matrix. For statistics, data from all triplicate measurements for both biosimilars were combined. A modified, asymmetric Gaussian curve was fitted to guide the eye (dotted curve). The median of the distribution is indicated by the vertical dashed, colored line. **b**, Example data for *δ*_ext_ > *δ*_int_ (left panel) and *δ*_ext_ < *δ*_int_ (right panel). For each treatment condition, i.e. controls or dilution series, a set of four cell culture wells was prepared (A-B; inter-well variability), for each of which four spots (i-iv; spot-to-spot variability) were measured on a MALDI target plate.

For both Rituximab batches (#1 and #2), the inter-well variability was higher than the spot-to-spot variability 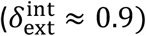 in case of the ATP readout. For the GSH readout both sources of variation were found to be similar 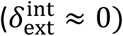. This discrepancy may be related to a more homogeneous matrix deposition process for the ATP readout (DHB, spray-coating) and thus a more robust spot-to-spot measurement. Alternatively, it may indicate partial ATP degradation during sample preparation leading to increased inter-well variability. In general, our data suggests that evaluation of 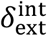 may help to identify sources of variation during assay development.

## Summary and Conclusion

In this study, we present the first proof-of-concept of the development of a quantitative MALDI-MS based cell bioassay for label-free assessment of therapeutic antibody potency. Relevant mAb concentration-dependent CDC mass markers were found in the low mass range at both polarities (metabolic and lipid mass range), and two metabolic compounds, ATP and GSH, were identified and validated as robust concentration-response markers in an untargeted approach. Obtained pEC_50_ values for rituximab CDC activity measured by MS compare well with the values obtained from luminescence-based readout method. In addition, the MALDI-TOF CDC bioassay provided comparable potency results for three different commercial antibody batches with suitable accuracy, variability and assay performance. Further improvements for use in HTS applications could be envisioned by automation of sample preparation steps like cell washing and incubation.

Regarding the MALDI-MS markers observed, the label-free detection of ATP exhibits a strongly correlation with, and serves to validate, the principle of the luminescence-based method, which quantifies ATP levels in surviving cells utilizing a luciferase-based reagent kit. GSH, on the other hand, plays a crucial role in the cells’ antioxidant defense and cell death^32^. While a full mechanistic understanding of these abundant markers in the CDC reaction are beyond the scope of this study, it demonstrates the ability of MALDI-TOF MS to monitor multiple markers based on their concentration response characteristics, enabling a multiplexing bioanalysis in a label-free setup. As presented in this study, the application of advanced MS technologies, such as TIMS TOF functionality, will further help to increase selectivity and identification of additional markers for future assay developments.

There is a significant interest in accelerating biopharmaceutical development by leveraging automation and high-throughput analytical technologies. Thus, functional bioassays using whole-cell MALDI-MS could provide a fast and cost-effective way to characterize the potency of new biotherapeutics biosimilars. Altogether, we envision high-throughput, label-free MALDI-MS cellular bioassays as a novel enabling tool for screening and characterization of complex and new modality biologics such as multi-specific antibodies or non-mAb therapeutics.

## Materials and Methods

### Materials and Cells

Acetonitrile (cat#34967) and methanol (cat#34966) were from Honeywell Specialty Chemicals Seelze GmbH (Seelze, Germany). RPMI-1640 with stable glutamine (cat#L0498-500), ESI-L Low concentration tuning mix (cat#HEWLG1969-85000), ammonium formate (cat#84884.180), isopropanol (cat#84881.320), 96-well tissue culture plates (cat#734-2327), Axygen 12-Channel Through (cat#732-1390) and reagent reservoirs (cat#89094-666 and cat#89094-668) were obtained from VWR International GmbH (Bruchsal, Germany). Trifluoracetic acid (cat# 1.08262.0025) and Triton X-100 (cat#1.12298.0101) were from Merck KGaA (Darmstadt, Germany), and N-(1-Naphthyl)ethylene diamine dihydrochloride (NEDC) (cat#222488-5G) and penicillin/streptomycin (PEN/Strep; cat# P0781) were from Sigma-Aldrich Chemie GmbH (Taufkirchen, Germany). Stable isotope-labelled ^13^C^1015^N_5_ -ATP (cat#CNLM-4265-CA-20) as internal standard was procured from Euriso-Top GmbH (Saarbrücken, Germany), and Raji cells (cat#CCL-86, ATCC) were purchased from LGC Standards GmbH (Wesel, Germany). 2,5-Dihydroxybenzoic acid (DHB) (cat#A11549), heat-inactivated fetal bovine serum (cat#15975161), Costar white microplates 3693 (cat# 10174381) were from Fisher Scientific GmbH (Schwerte, Germany). CellTiter-Glo reagent (cat#G7572) was obtained from Promega GmbH (Walldorf, Germany) and stored in aliquots at -20 °C after preparation. Rabbit complement, unadsorbed (cat# 239-31060-1) was from Gentaur GmbH (Aachen, Germany), and 96-well clear microplates (cat#675161) were from Greiner Bio-One GmbH (Frickenhausen, Germany). Deep-Well plates (cat#0030501306) were from Eppendorf Liquid Handling GmbH (Hamburg, Germany). Anchor chip MALDI target plates (MTP) (cat#8280790) and polished steel MTPs (cat#8280781; Bruker Daltonics GmbH, Bremen, Germany) were used. Rituximab drug product batches (MabThera®) #1, #2 and #ref were from Roche.

### Cell Culture

Raji Cells were suspension-cultured in RPMI-1640 medium supplemented with stable glutamine, 10% FCS, 1% PEN/Strep. Cells were maintained by seeding 2.5*10^5^ cells/mL in T75 flasks every two days. For assays, cells were harvested by centrifugation, supernatant was removed and cells resuspended in warm medium at 5·10^6^ cells/mL. Cells were transferred to a reservoir, and 50 µL of the cell suspension was transferred per well into a 96-well flat-bottom plate using an 8-channel pipette in dispensing mode shortly (< 30 min) before this “cell plate” was used in the assay.

### Complement-dependent Cytotoxicity (CDC) Assay

MS-based CDC assay development was based on a published procedure^15^. Complement was thawed on ice and diluted 1:2 with ice-cold assay medium. Assay medium, diluted complement and 2% Triton X-100 in assay medium were transferred into a reservoir, a 96-well half-area microplate containing 0.36 mg/mL antibody. This “complement plate” and an empty half-area microplate were placed in a CyBio FeliX liquid handling device equipped with an 8-channel adapter (Analytik Jena GmbH, Jena, Germany) for preparation of the “sample plate”. This contained an antibody dilution series for final assay concentrations of 0.83 µM, 0.21 µM, 0.05 µM, 0.01 µM, 6.5 nM, 3.3 nM, 0.8 nM, 0.2 nM and 0.05 nM. 50 µL each from the sample and the complement plate were transferred to the cell plate, which was henceforth referred to as the “assay plate”. It contained a “blank” (non-treated cells) as well as negative (treated with complement only) and positive controls (treated with Triton X-100 to cause cell death). The assay plate was then incubated at 37 °C, 5% CO_2_ for 2h in a HERAcell 150 cell culture incubator (Thermo Electron LED GmbH, Langenselbold, Germany) and then placed in the liquid handling device.

### Luminescence assay

Cells were suspended, and 6 µL of the cell solution was transferred into a white half-area plate containing 50 µL CellTiter Glo reagent for reference luminescence measurement. It was shaken for 10 min and measured on a POLARstar Omega microplate reader (BMG Labtech GmbH, Offenburg, Germany) with a 1 s measurement interval per well.

### MALDI MS assay plate preparation

Meanwhile the assay plate was centrifuged (MEGA STAR 4.0R, VWR International BVBA, Leuven, Belgium) at 500 g, 4 °C for 5 min, and the supernatant was removed. Ice-cold washing solution (150 µL of 150 mM ammonium formate, pH 7.6) was added to each well, and the centrifugation step was repeated. Washing solution was aspirated, the cell plate was flash-frozen in liquid nitrogen and stored at -80 °C.

### MALDI MS assay

The assay plate was placed in the CyBio FeliX. Resuspension solvent (50% acetonitrile with 20 µM isotope-labelled ATP) was added, and cells were suspended by automated pipetting and shaking. Excess cell suspension was aspirated, and the cell suspension was spotted on an anchor chip 384 BC MTP and a polished steel MTP as quadruplicates. Spotting all samples of a 96-well plate on a 384-spot MTP as quadruplicates took about 10 s with a 96-tip adapter. Target plates were air-dried before matrix deposition. Then 1 µL of 7 mg/mL NEDC matrix dissolved in 70% methanol was deposited on each sample spot of the polished steel MTP. For matrix deposition by spotting, a half-area microplate was prepared containing 50 µL of the respective matrix solution. The CyBio FeliX aspirated excess matrix solution, waited for 5 seconds to saturate the gas phase, dispensed 1 µL, spotted one microliter on a waste plate and continued with matrix deposition, before the plate was air-dried. DHB matrix deposition followed a published protocol^18^. In brief, DHB was dissolved in 50% acetonitrile, 2.5% TFA at a concentration of 20 mg/mL and deposited on the anchor chip using a TM3 sprayer (HTX Technologies, Chapel Hill, USA) with the following settings: 60 °C nozzle temperature, four passes, 60 µL/min flowrate, 1000 mm/min velocity, 2 mm track spacing, 10 psi gas pressure and 40 mm nozzle height^19^. After drying, plates were measured in a rapifleX MALDI-ToF mass spectrometer using FlexControl Version 4.2 (Bruker Daltonics). Methods were calibrated in the 300-600 *m/z* range using red phosphor clusters^33^. MTPs were automatically measured in positive and negative ion modes. Two out of the three replicates were additionally measured in a MALDI-qToF mass spectrometer using timsControl Version 4.1.8 and calibration on tune-mix (timsTOF fleX, Bruker Daltonics) to use ion mobility for evaluation if identified markers contained any underlying molecular signal. For automated data acquisition in *screening / analyze* mode, timsTOF MALDI Pharma Pulse (MPP) 2023 software (Bruker) was used: Isotope-labelled ATP was set as internal standard (*m/z* 523.0217). Glutathione ([M-H]^-^; *m/z* 306.0765) was selected as target compound in negative ion mode in the *m/z* range 200-1300 Da for both timsOFF/QTOF mode and timsON measurements. The settings for timsON were: 1/K0 Start: 0.50 V·s/cm^2^, 1/K_0_ End: 1.00 V·s/cm^2^, Ramp time 60.0 ms. Accumulation time was 14.8 ms, duty cycle 24.67% and the Ramp time 15.36 Hz. Laser application *imaging 100 µm* with 148 shots at 10,000 Hz with 10 shots per raster position and random walk enabled over the complete sample with a spot diameter of 2000 µm were used. The transfer time was set to 80 µs and 8 µs pre-pulse storage. For the timsOFF method 1000 shots at 2000 Hz with 100 shots per raster position and random walk enabled over the complete sample with a spot diameter of 2000 µm were used. The transfer time was set to 50 µs and the pre-pulse storage to 7.0 µs.

### Data Analysis and Statistics

For identification of molecular markers, MS spectral data was analyzed using an in-house software tool^15^ with slight modifications. During data import, data smoothing (Savitzky-Golay) and baseline subtraction (Top-hat) were performed. The signal-to-noise threshold was 3 for peak detection. Spectra were total ion current (TIC) normalized and single-point recalibrated with a tolerance of 0.1 Da on the internal standard if present – alternatively on phosphatidylcholine 34:1 (*m/z* 760.585). Alignment tolerance was 0.01 Da and binning tolerance 200 ppm. Based on the log-2 fold-changes, candidate *m/z* were selected, and their curve shape and pEC_50_ value was determined. Peak identification was performed on a timsTOF flex MALDI mass spectrometer for two prominent *m/z* values with high log-2 fold-change and signal intensity. The peaks were isolated and fragmented with different fragmentation energies to achieve optimal fragment distribution. For evaluation of CDC assays based on identified markers, MALDI-TOF MS data was loaded into R for automatic extraction of *m/z* values. After setting the normalization method (internal standard for ATP, TIC for glutathione) the *m/z* value to be extracted (*m/z* 508.02 for ATP, *m/z* 306.075 for glutathione) was chosen.

Data acquired using timsTOF MALDI Pharma Pulse was exported in the .csv data format. TimsOFF spectra were evaluated based on glutathione peak intensity at *m/z* 306.075, while the peak area was used for timsON measurements. Graphpad prism 5.0 (GraphPad Software, Boston, USA) and a 4-parameter non-linear regression (curve fit) was used to fit the data. Calculated pEC_50_ values were used for comparison between measurements. To assess the quality of the MALDI MS-based CDC assay, the Z’ and V’ factors were used. The Z’ factor was calculated according to Zhang et al.^29^.

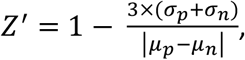

where *μ*_*p*_ and *μ*_*n*_ is the mean value of the positive and negative control, respectively. The V’ values were determined according to Bray et al.^30^.

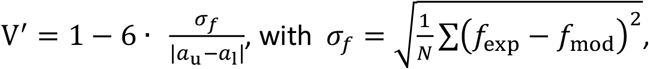

where *σf* is the standard deviation of the residuals of the 4-parameter non-linear regression model f calculated from the experimental (exp) data and the fit (mod). *a*_u_ and *a*_l_ are the values of the upper (u) and lower (l) asymptotes from the model fit.

The ratio of inter-well and spot-to-spot variance is determined by introducing a quality measure 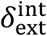 defined as *δ*_ext_/*δ*_int_ − 1 for *δ*_ext_ > *δ*_int_ and −(*δ*_int_/*δ*_ext_ − 1) for *δ*_ext_ < *δ*_int_. *δ*_ext_ and *δ*_int_ are given as 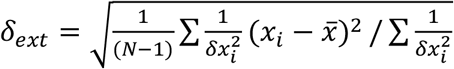 and 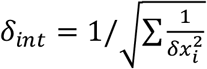, where *x*_*i*_ and *δ*_*xi*_ are the mean values and related uncertainties (standard deviation) obtained from the repeated measurements^34^. 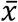 denotes the mean value and *N* is the number of data points. Within this work, in total 16 data points per treatment condition were measured, i.e. 4 technical replicates per *well* and *spot* each.

## Author Contributions

The study was conceived by C.H. and C.G. S.S. designed experiments, coordinated the work and performed extensive data analysis. A.G. developed the MALDI MS assay, designed most experiments and performed all experiments including cell culture, MS measurements and lab automation. T.E. contributed bioinformatics for the untargeted identification of biomarkers. B.F. developed assay automation protocols. L.P and M.V. evaluated luminescence data, suggested experiments and provided materials and samples. C.G. coordinated the project, contributed to the evaluation of the results, and edited the manuscript. C.H. supervised the overall work, evaluated the results and provided infrastructure. A.G., S.S. and C.H. wrote the manuscript. All authors have read and agreed to the published version of the manuscript.

## Acknowledgments

We thank Lars Gruber for fruitful discussion on fragmentation analysis and ion mobility spectrometry and Dariusz Janecki for helpful discussions about MS. This work was mainly funded by Novartis. Acquisition of the rapifleX was supported by the Hector Foundation II to C.H., acquisition of the timsTOF flex by the German Federal Ministry of Research (BMBF) to C.H. as part of the MSCoreSys SMART-CARE project (Project 16LW0238). C.H. acknowledges the support by the BMBF (German Federal Ministry of Research) as part of the Innovation Partnership “Multimodal Analytics and Intelligent Sensorics for the Health Industry” (M^2^Aind), projects “Drugs4Future” (grant 12FH8I05IA) and “DrugsData” (grant 13FH8I09IA) and by the Ministerium für Wissenschaft, Forschung and Kunst (MWK) Baden-Württemberg by the Mittelbauprogramm.

## Data Availability Statement

The datasets generated during and/or analysed during the current study are available from the corresponding author on reasonable request.

## Competing interests

C.G., V.M. and L.P. were employed by Novartis at the time of the study. Novartis provided materials (e.g., antibodies) and funded the research. S.S., A.G., T.E., B.F. and C.H. declare no other competing interests.

## References

1 Kesik-Brodacka, M. Progress in biopharmaceutical development. Biotechnol Appl Biochem 65, 306–322, doi:10.1002/bab.1617 (2018).

2 Dash, R., Singh, S. K., Chirmule, N. & Rathore, A. S. Assessment of Functional Characterization and Comparability of Biotherapeutics: a Review. AAPS J 24, 15, doi:10.1208/s12248-021-00671-0 (2021).

3 Berkowitz, S. A., Engen, J. R., Mazzeo, J. R. & Jones, G. B. Analytical tools for characterizing biopharmaceuticals and the implications for biosimilars. Nat Rev Drug Discov 11, 527–540, doi:10.1038/nrd3746 (2012).

4 Khetan, R. et al. Current advances in biopharmaceutical informatics: guidelines, impact and challenges in the computational developability assessment of antibody therapeutics. MAbs 14, 2020082, doi:10.1080/19420862.2021.2020082 (2022).

5 Lei, Y., Yong, Z. & Junzhi, W. Development and application of potency assays based on genetically modified cells for biological products. J Pharm Biomed Anal 230, 115397, doi:10.1016/j.jpba.2023.115397 (2023).

6 Wieckowski, S., Avenal, C., Orjalo, A. V., Jr., Gygax, D. & Cymer, F. Toward a Better Understanding of Bioassays for the Development of Biopharmaceuticals by Exploring the Structure-Antibody-Dependent Cellular Cytotoxicity Relationship in Human Primary Cells. Front Immunol 11, 552596, doi:10.3389/fimmu.2020.552596 (2020).

7 White, J. R. et al. Best practices in bioassay development to support registration of biopharmaceuticals. Biotechniques 67, 126–137, doi:10.2144/btn-2019-0031 (2019).

8 Nimmerjahn, F. & Ravetch, J. V. Fcgamma receptors as regulators of immune responses. Nat Rev Immunol 8, 34–47, doi:10.1038/nri2206 (2008).

9 Rossignol, A., Bonnaudet, V., Clemenceau, B., Vie, H. & Bretaudeau, L. A high-performance, non-radioactive potency assay for measuring cytotoxicity: A full substitute of the chromium-release assay targeting the regulatory-compliance objective. MAbs 9, 521–535, doi:10.1080/19420862.2017.1286435 (2017).

10 Chand, S. et al. A reliable assay for ensuring the biological activity of anti T lymphocyte immunoglobulin as an alternate to compendial flow cytometry method. Biologicals 65, 33–38, doi:10.1016/j.biologicals.2020.01.002 (2020).

11 Salinas-Jazmin, N., Medina-Rivero, E. & Velasco-Velazquez, M. A. Bioassays for the Evaluation of Target Neutralization and Complement-Dependent Cytotoxicity (CDC) of Therapeutic Antibodies. Methods Mol Biol 2313, 281–294, doi:10.1007/978-1-0716-1450-1_17 (2022).

12 Belov, A. M. et al. Acoustic Mist Ionization-Mass Spectrometry: A Comparison to Conventional High-Throughput Screening and Compound Profiling Platforms. Anal Chem 92, 13847–13854, doi:10.1021/acs.analchem.0c02508 (2020).

13 Simon, R. P. et al. Acoustic Ejection Mass Spectrometry: A Fully Automatable Technology for High-Throughput Screening in Drug Discovery. SLAS Discov 26, 961–973, doi:10.1177/24725552211028135 (2021).

14 Duenas, M. E. et al. Advances in high-throughput mass spectrometry in drug discovery. EMBO Mol Med 15, e14850, doi:10.15252/emmm.202114850 (2023).

15 Unger, M. S., Blank, M., Enzlein, T. & Hopf, C. Label-free cell assays to determine compound uptake or drug action using MALDI-TOF mass spectrometry. Nat Protoc 16, 5533–5558, doi:10.1038/s41596-021-00624-z (2021).

16 Pu, F. et al. New Platform for Label-Free, Proximal Cellular Pharmacodynamic Assays: Identification of Glutaminase Inhibitors Using Infrared Matrix-Assisted Laser Desorption Electrospray Ionization Mass Spectrometry. ACS Chem Biol 18, 942–948, doi:10.1021/acschembio.3c00087 (2023).

17 Munteanu, B. et al. Label-free in situ monitoring of histone deacetylase drug target engagement by matrix-assisted laser desorption ionization-mass spectrometry biotyping and imaging. Anal Chem 86, 4642–4647, doi:10.1021/ac500038j (2014).

18 Weigt, D., Sammour, D. A., Ulrich, T., Munteanu, B. & Hopf, C. Automated analysis of lipid drugresponse markers by combined fast and high-resolution whole cell MALDI mass spectrometry biotyping. Sci Rep 8, 11260, doi:10.1038/s41598-018-29677-z (2018).

19 Weigt, D. et al. Mechanistic MALDI-TOF Cell-Based Assay for the Discovery of Potent and Specific Fatty Acid Synthase Inhibitors. Cell Chem Biol 26, 1322–1331 e1324, doi:10.1016/j.chembiol.2019.06.004 (2019).

20 Delannoy, C. P. et al. High-Throughput Quantitative Screening of Glucose-Stimulated Insulin Secretion and Insulin Content Using Automated MALDI-TOF Mass Spectrometry. Cells 12, doi:10.3390/cells12060849 (2023).

21 Heap, R. E., Segarra-Fas, A., Blain, A. P., Findlay, G. M. & Trost, M. Profiling embryonic stem cell differentiation by MALDI TOF mass spectrometry: development of a reproducible and robust sample preparation workflow. Analyst 144, 6371–6381, doi:10.1039/c9an00771g (2019).

22 Unger, M. S. et al. Direct Automated MALDI Mass Spectrometry Analysis of Cellular Transporter Function: Inhibition of OATP2B1 Uptake by 294 Drugs. Anal Chem 92, 11851–11859, doi:10.1021/acs.analchem.0c02186 (2020).

23 Prior, S. et al. International standards for monoclonal antibodies to support pre-and postmarketing product consistency: Evaluation of a candidate international standard for the bioactivities of rituximab. MAbs 10, 129–142, doi:10.1080/19420862.2017.1386824 (2018).

24 Kennedy, L., Sandhu, J. K., Harper, M. E. & Cuperlovic-Culf, M. Role of Glutathione in Cancer: From Mechanisms to Therapies. Biomolecules 10, doi:10.3390/biom10101429 (2020).

25 Greiner, J. V. & Glonek, T. Intracellular ATP Concentration and Implication for Cellular Evolution. Biology (Basel) 10, doi:10.3390/biology10111166 (2021).

26 Flood, D., Lee, E. S. & Taylor, C. T. Intracellular energy production and distribution in hypoxia. J Biol Chem 299, 105103, doi:10.1016/j.jbc.2023.105103 (2023).

27 Plante, P. L. et al. Predicting Ion Mobility Collision Cross-Sections Using a Deep Neural Network: DeepCCS. Anal Chem 91, 5191–5199, doi:10.1021/acs.analchem.8b05821 (2019).

28 Zhang, H. et al. AllCCS2: Curation of Ion Mobility Collision Cross-Section Atlas for Small Molecules Using Comprehensive Molecular Representations. Anal Chem 95, 13913–13921, doi:10.1021/acs.analchem.3c02267 (2023).

29 Zhang, J. H., Chung, T. D. & Oldenburg, K. R. A Simple Statistical Parameter for Use in Evaluation and Validation of High Throughput Screening Assays. J Biomol Screen 4, 67–73, doi:10.1177/108705719900400206 (1999).

30 Bray, M. A. & Carpenter, A. in Assay Guidance Manual (eds S. Markossian et al.) (2004).

31 Roux, C. et al. Data analysis of Q-value measurements for double-electron capture with SHIPTRAP. Eur. Phys. J. D 67, 146, 10.1140/epjd/e2013-40110-x (2013).

32 Franco, R. & Cidlowski, J. A. Apoptosis and glutathione: beyond an antioxidant. Cell Death Differ 16, 1303–1314, doi:10.1038/cdd.2009.107 (2009).

33 Sladkova, K., Houska, J. & Havel, J. Laser desorption ionization of red phosphorus clusters and their use for mass calibration in time-of-flight mass spectrometry. Rapid Commun Mass Spectrom 23, 3114–3118, doi:10.1002/rcm.4230 (2009).

34 Birge, R. T. The Calculation of Errors by the Method of Least Squares. Phys. Rev. 40, 10.1103/PhysRev.40.207 (1932).

