## Supplementary information for "Label-free Assessment of Complement-Dependent Cytotoxicity of Therapeutic Antibodies via a Whole-Cell MALDI Mass Spectrometry Bioassay"


| | | |
|---|---|---|
| SI Figure 1 (Fig. S1) | Assay characteristics and optimized assay conditions. | S-2 |
| SI Figure 2 (Fig. S2) | Concentration-responses of an identified CDC- and Rituximab response marker | S-3 |
| SI Figure 3 (Fig. S3) | Identification of adenine triphosphate as a CDC response markers | S-3 |
| SI Figure 4 (Fig. S4) | Individual MALDI-TOF MS drug response curves for ATP for three different Rituximab drug product samples, #1, #2 and #ref. | S-4 |
| SI Figure 5 (Fig. S5) | Individual MALDI-TOF MS drug response curves for GSH for three different Rituximab drug product samples, #1, #2 and #ref. | S-5 |
| SI Figure 6 (Fig. S6) | Representative drug response curves for glutathione for different measurement devices and modes of operation for GSH | S-6 |
| SI Table 1 (Tab. S1) | $pEC_{50}$ values for 3 different Rituximab drug product samples and readouts | S-7 |

#### Supplementary Figures

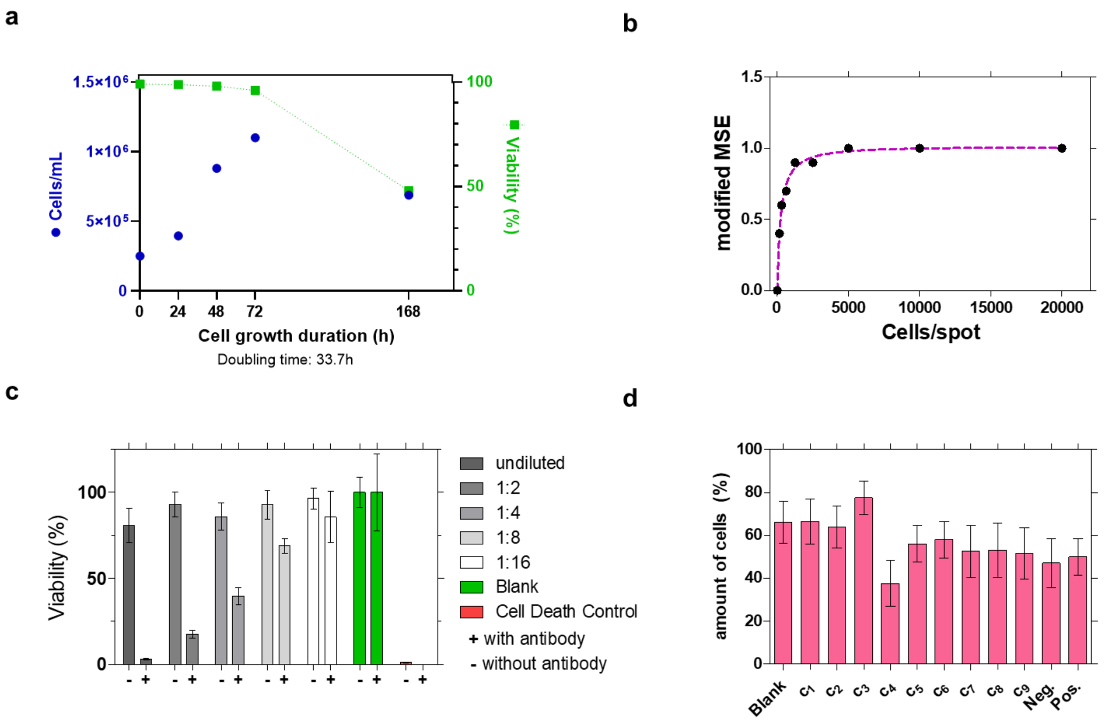


**Figure S1. Assay characteristics and optimized assay conditions. a**, Cell growth and viability of Raji cells. Doubling times and maximum cell density were in good agreement with information from the supplier. **b**, Evaluation of the modified Matrix Suppression Effect (MSE) revealed an optimal cell number of 5000 cells per spot. The dashed fit of a saturation curve was included to guide the eye. **c**, Cell viability in percent of blank (non-treated cells) suggested an optimal complement dilution of 1:2. A Rituximab concentration of 52 nM was used in the assay. **d**, Relative number of cells, as percent of the number of seeded cells after washing (step 3 in **Figure 1**) for different treatment conditions, i.e. blank, antibody concentrations $c_1$ = 0.83 µM to $c_9$ = 0.05 nM, negative (neg.) and positive (pos.) controls.

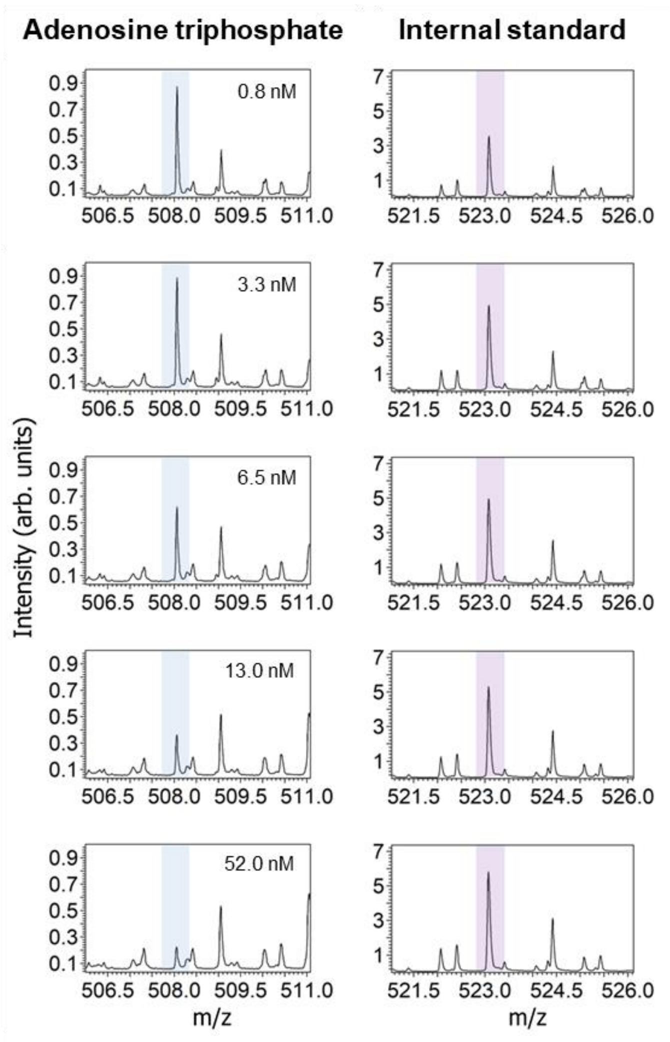


**Figure S2. Concentration-responses of an identified CDC- and Rituximab response marker (adenosine triphosphate) and its $^{13}C_{10}{}^{15}N_5$ stable isotope labeled internal standard.** Spectra are depicted for concentrations of the antibody ranging from 0.8 nM to 52.0 nM.

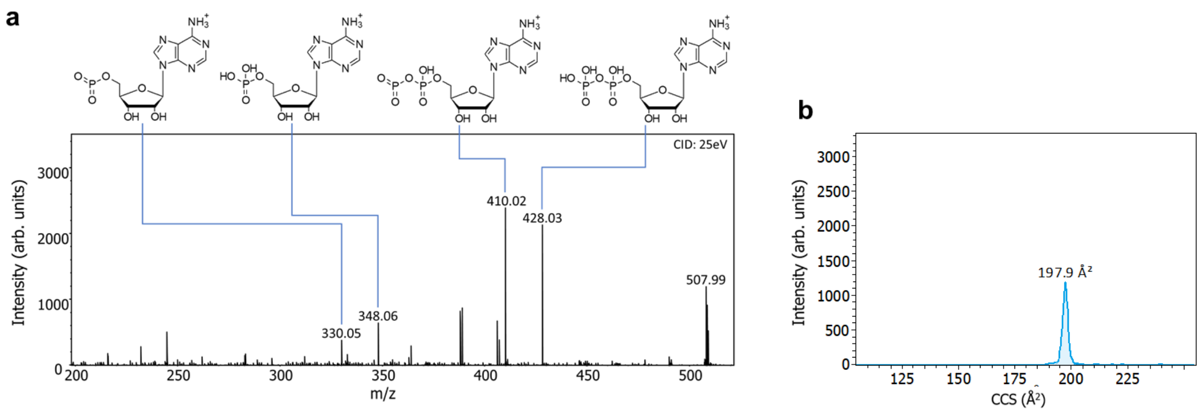


**Figure S3. Identification of adenosine triphosphate as a CDC response markers. a**, MALDI-MS/MS spectra of *m/z* 508.00 display unique fragments of ATP (theoretical m/z 508.0030; DHB matrix) as indicated by corresponding fragment structures. Measurements were performed on the timsTOF flex mass spectrometer. Ion mobilogram for ATP for experimental deduction of collisional cross section (CCS) of 198 ± 2 Å$^2$ for ATP. For these response markers there was no indication of interfering compounds.

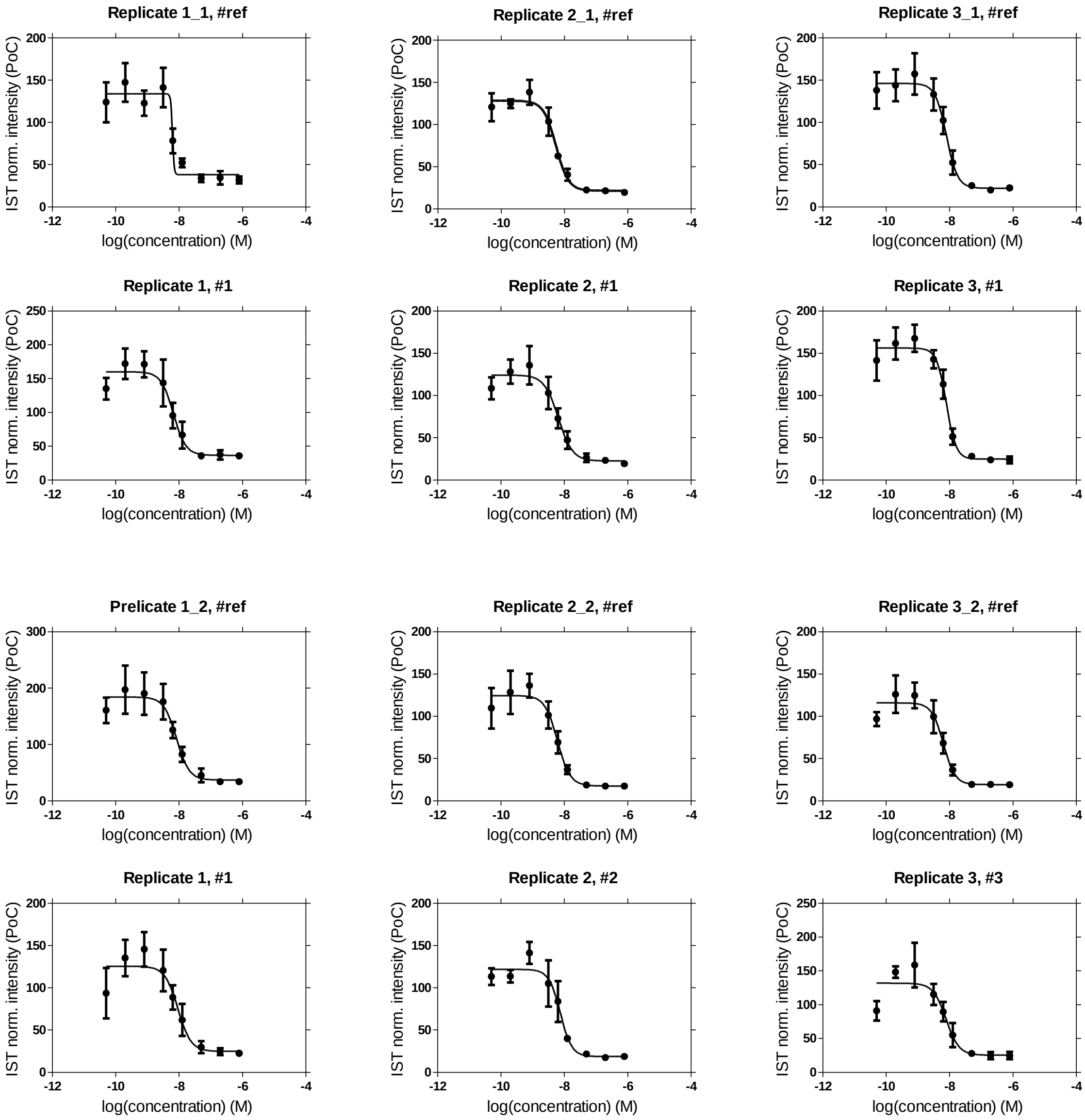


**Figure S4. Individual MALDI-TOF MS drug response curves for ATP for three different Rituximab drug product samples, #1, #2 and #ref**. Uncertainties are given as standard deviation

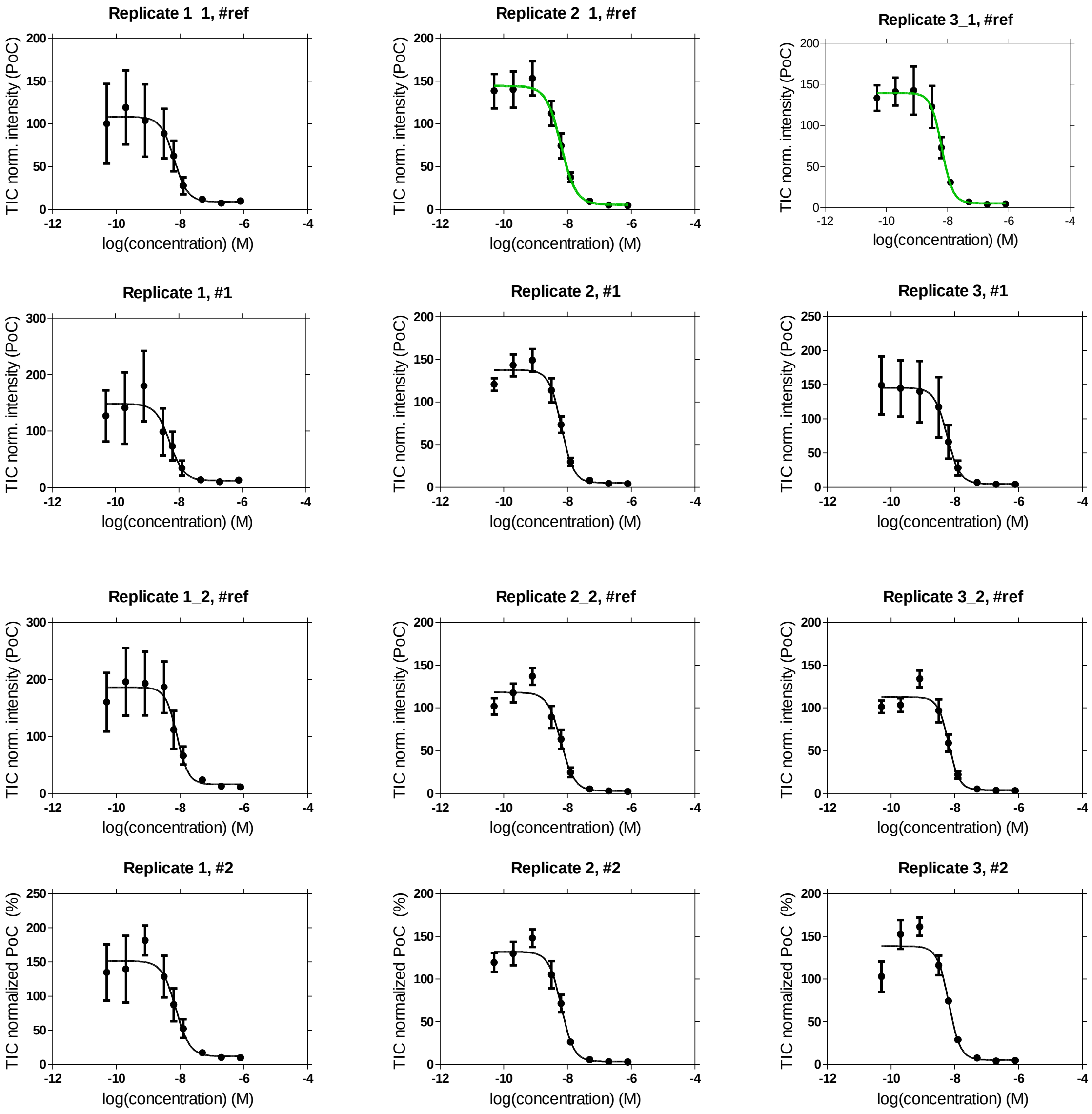


**Figure S5. Individual MALDI-TOF MS drug response curves for GSH for three different Rituximab drug product samples, #1, #2 and #ref.** Uncertainties are given as standard deviation.

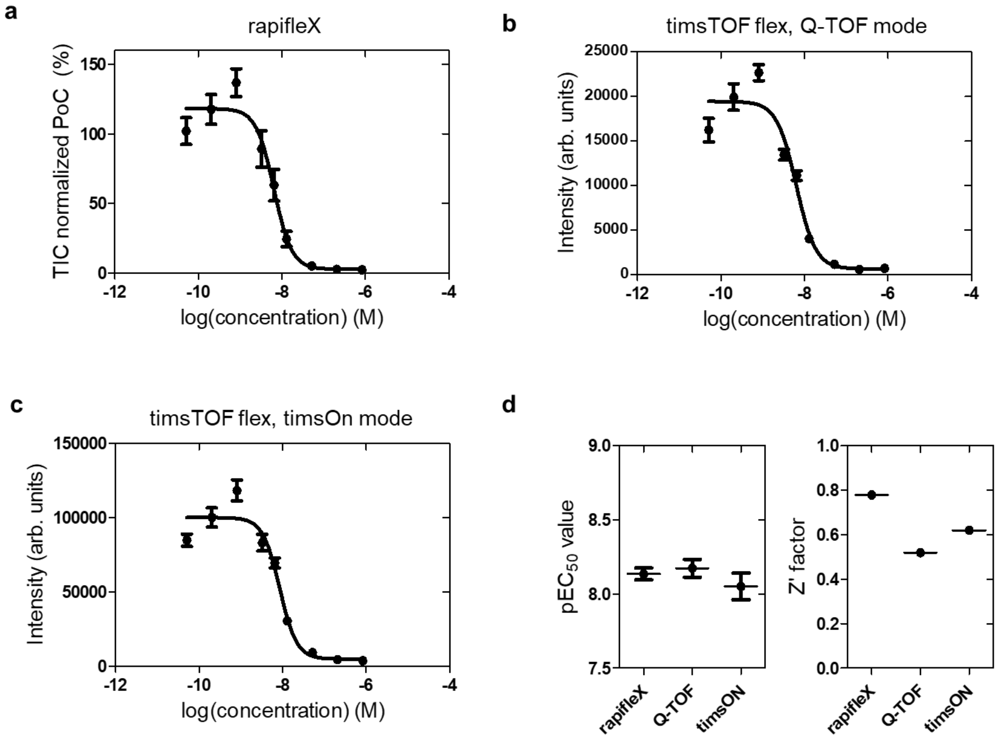


**Figure S6**: **Representative drug response curves for glutathione for different measurement devices and modes of operation for GSH. a** rapifleX MALDI-TOF data, **b** timsTOF flex data in Q-TOF mode and **c** timsTOF flex data in timsOn mode. **d**, $pEC_{50}$ values and Z' factors of the corresponding assays of the reference antibody (#ref). $pEC_{50}$ values were 8.14 ± 0.04 , 8.17 ± 0.06 and 8.05 ± 0.09 for the three instruments, respectively, determined for one replicate. Values for the Z' factor were 0.78, 0.52 and 0.62, respectively.

### Supplementary Tables

**Table S1**: **CDC pEC$_{50}$ values for different Rituximab drug product batch samples (#ref, #1, #2) and readouts.** [1] Same antibody, which serves as reference in the comparability study. [2]Data shown in Fig. 3b.

| | #ref[1] | #1 | #ref[1] | #1 | |
|---|---|---|---|---|---|
| pEC$_{50}$, rep. 1 | 8.21 ± 0.05 | 8.18 ± 0.03 | 8.08 ± 0.03[2] | 8.03 ± 0.04 | **MALDI MS, ATP** |
| pEC$_{50}$, rep. 2 | 8.27 ± 0.02 | 8.19 ± 0.03 | 8.22 ± 0.03[2] | 8.12 ± 0.03 | |
| pEC$_{50}$, rep. 3 | 8.10 ± 0.02 | 8.10 ± 0.02 | 8.19 ± 0.03[2] | 8.11 ± 0.04 | |
| pEC$_{50}$, mean | **8.18 ± 0.06** | **8.14 ± 0.03** | **8.16 ± 0.04[2]** | **8.09 ± 0.03** | |
| pEC$_{50}$, rep. 1 | 8.18 ± 0.05 | 8.28 ± 0.06 | 8.10 ± 0.04[2] | 8.13 ± 0.04 | **MALDI MS, GSH** |
| pEC$_{50}$, rep. 2 | 8.20 ± 0.02 | 8.19 ± 0.02 | 8.20 ± 0.02[2] | 8.19 ± 0.02 | |
| pEC$_{50}$, rep. 3 | 8.18 ± 0.02 | 8.24 ± 0.05 | 8.19 ± 0.02[2] | 8.19 ± 0.03 | |
| pEC$_{50}$, mean | **8.19 ± 0.01** | **8.20 ± 0.02** | **8.18 ± 0.02** | **8.18 ± 0.01** | |
| pEC$_{50}$, rep. 1 | 8.28 ± 0.04 | 8.33 ± 0.04 | 8.24 ± 0.03[2] | 8.24 ± 0.05 | **Luminenscence** |
| pEC$_{50}$, rep. 2 | 8.15 ± 0.03 | 8.18 ± 0.03 | 8.15 ± 0.04[2] | 8.14 ± 0.04 | |
| pEC$_{50}$, rep. 3 | 8.14 ± 0.04 | 8.15 ± 0.04 | 8.17 ± 0.04[2] | 8.19 ± 0.04 | |
| pEC$_{50}$, mean | **8.18 ± 0.04** | **8.21 ± 0.05** | **8.20 ± 0.03** | **8.18 ± 0.03** | |
| | **8.19 ± 0.01** | **8.18 ± 0.02** | **8.19 ± 0.01** | **8.17 ± 0.01** | |
